# Microcolonies: a novel morphological form of pathogenic *Mycoplasma* spp

**DOI:** 10.1101/535559

**Authors:** Irina V. Rakovskaya, Svetlana A. Ermolaeva, Galina A. Levina, Olga I. Barkhatova, Andrey Ya. Mukhachev, Svetlana G. Andreevskaya, Vladimir G. Zhukhovitsky, Luisa G. Gorina, Galina G. Miller, Elena V. Sysolyatina

## Abstract

The work describes a novel morphological form found in 5 species of *Mollicutes*: *Mycoplasma hominis, M. fermentans, M. gallisepticum, M. pneumoniae, Acholeplasma laidlawii*. The form, which is referred to as microcolonies (MCs) in contrast to typical colonies (TCs), was characterized by tiny propeller-shaped colonies formed by rod-like cells tightly packed in parallel rows. MCs were observed within routinely cultivated type strain cultures of the listed species, and rod-like cells could be seen ewith SEM within TCs of the “fried-egg” type. Pure MC cultures were obtained by treatment of Mycoplasma cultures with hyperimmune serum, antibiotics or argon non-thermal plasma. Appearance of MCs was observed 7-12 days post plating while TCs appeared 24-48 h post plating. MCs derived from the *M. hominis* type strain H-34 were characterized in details. MCs did not differ from the parental culture in the MLST, direct fluorescent and epifluorescent tests and Western-blotting with a monospecific camel-derived nanoantibody aMh-FcG2a. Meanwhile, MCs derived from this strain and other listed species were resistant to at least 9 antibiotics and did not hydrolyze arginine and/or glucose in standard bacteriological tests. MC cultures that belonged to *M. hominis* (n=70), *M. pneumoniae* (n=2), *M. fermentans* (n=2), *Mycoplasma* spp (n=5) were isolated from clinical samples of serum, synovial liquid and urina of patients with inflammatory urogenital tract diseases, asthma, arthritis. The reported MCs might be similar to “small colony variants” (SCVs) described in other bacterial species. However, in contrast to SCVs, MCs have never reverted to TCs. Multiple consecutive re-plating steps (for up to 3 years) were not sufficient to provide appearance of TCs within a pure MC culture. An unknown role of MCs in infection pathology along with their prominent antibiotic resistance makes them a challenge for the future studies of *Mollicutes*.

**Author summary:** Here we demonstrated that Mycoplasma species form small size colonies (referred to as minicolonies, MCs). MC size is ten times less than the size of typical Mycoplasma colonies (TCs). MCs are very slow growing: it was required 9-10 days for MCs to form in contrast to 24-72 h required for TCs to form. The presents a system of evidences that MCs are formed by the same species as TCs, which they have been obtained from. Pure culture of MCs might be obtained from TC cultures by treatment with the hyperimmune serum, antibiotic and non-thermal gas plasma. MCs of all species were resistant to antibiotics effective against TCs. MCs did not hydrolyze arginine and glucose in standard bacteriological tests. MCs of different Mycoplasma species were isolated from clinical samples of sera, urea and synovial fluids from patients with urolithiasis, rheumatoid arthritis and asthma. MCs never have reverted to TCs even after three years passing. A role of MCs in infectious pathology has not been established yet. Nevertheless, ability to persist in the human body and extreme antibiotic resistance make MCs to be a challenge for the future research.

## Introduction

*Mycoplasma spp* belong to the *Mollicutes* class that united cell wall lacking bacteria most of which are membrane parasites [1–3]. *Mycoplasma spp* often colonize mucosal surfaces of the urogenital and respiratory tracts in a nonvirulent manner and spread over the organism to other organs. Being opportunistic bacteria, *Mycoplasmas* may cause postoperative, postpartum and posttraumatic infections in various systems. *Mycoplasma* spp are etiologic agents of pneumonias, non-gonococcal urethritis, endometritis, and chorioamnionitis, and infertility in both immunosuppressed and immunocompetent individuals [4–8]. Some of *Mycoplasma* play role in development of autoimmune diseases such as asthma [9–11].

*Mycoplasma* spp when they are grown on the agar form colonies with the dense partly immersed in the agar center and delicate periphery, the morphology is known as “fried-egg”[12–14]. These colonies that further we will call typical colonies (TCs) grow up to the full size within 48-72 h and have a diameter ranges from about 75 to 800 µm [15]. *Mycoplasma* colonies might be seen with the naked eye as very small dots but usually TCs are observed with light microscopy. Multiple studies demonstrated that *Mycoplasma* colonies consist of variable elements including spheres and strings of different electronic density that lay separately or form structures of a higher order [16,17]. *Mycoplasma* cells range in size from 200 to 800 nm with the smallest cells of about 100-125 nm [2,17]. Under laboratory conditions, generation rates are about 1,6-2 h for colony formation to take about 48-72 h [12,17,18]. In certain cases, e.g. for isolates obtained from clinical samples, these time intervals increase.

Dealing with an analysis of clinical samples for *Mycoplasma* spp for many years, we combine bacteriological methods with immunological methods and PCR. When clinical samples are PCR-positive, we perform a bacteriological assay. It is known that usually serum is free of Mycoplasma spp, and we failed to observed TCs in PCR-positive samples within standard cultivation time. However, prolonging incubation time of immune- and/or PCR-positive serum samples up to 7-9 days or sometimes 12 days, we revealed very small colonies of non-round morphology. Such colonies were highly likely to defects on the agar surface. Meantime, they were not. These colonies, which are referred below as minicolonies (MCs) (i) could be re-plated from 2-10 times and up to 3 years; (ii) gave a positive signal in PCR in the absence of visible TCs and interacted with *Mycoplasma* specific antibodies; (iii) could be washed out from the agar surface to get samples for checking out in Western blotting (see below), proteomic (see elsewhere) and genomic (see elsewhere) assays. Up to our knowledge, such type colonies have not been described yet. This paper is devoted to characterization of MCs formed by *Mycoplasma* spp.

## Results

### First evidences of minicolonies (MCs) from clinical samples

It is well known that *Mycoplasma* spp are rarely isolated from clinical blood/serum samples. Meanwhile, we had often observed positive signals when applied PCR to test serum samples. As these results suggested a *M. hominis* infection, we plated serum samples according to the procedure but we have never observed typical *M. hominis* colonies (TCs) in these conditions. However, prolonged incubation time up to 7-9 and sometimes 12 days resulted in appearance of very small colonies with non-typical morphology that were highly likely to artefacts appearing on the agar surface (Fig. 1, A and B). We referred to this type colonies as minicolonies (MCs).

**Fig. 1.**
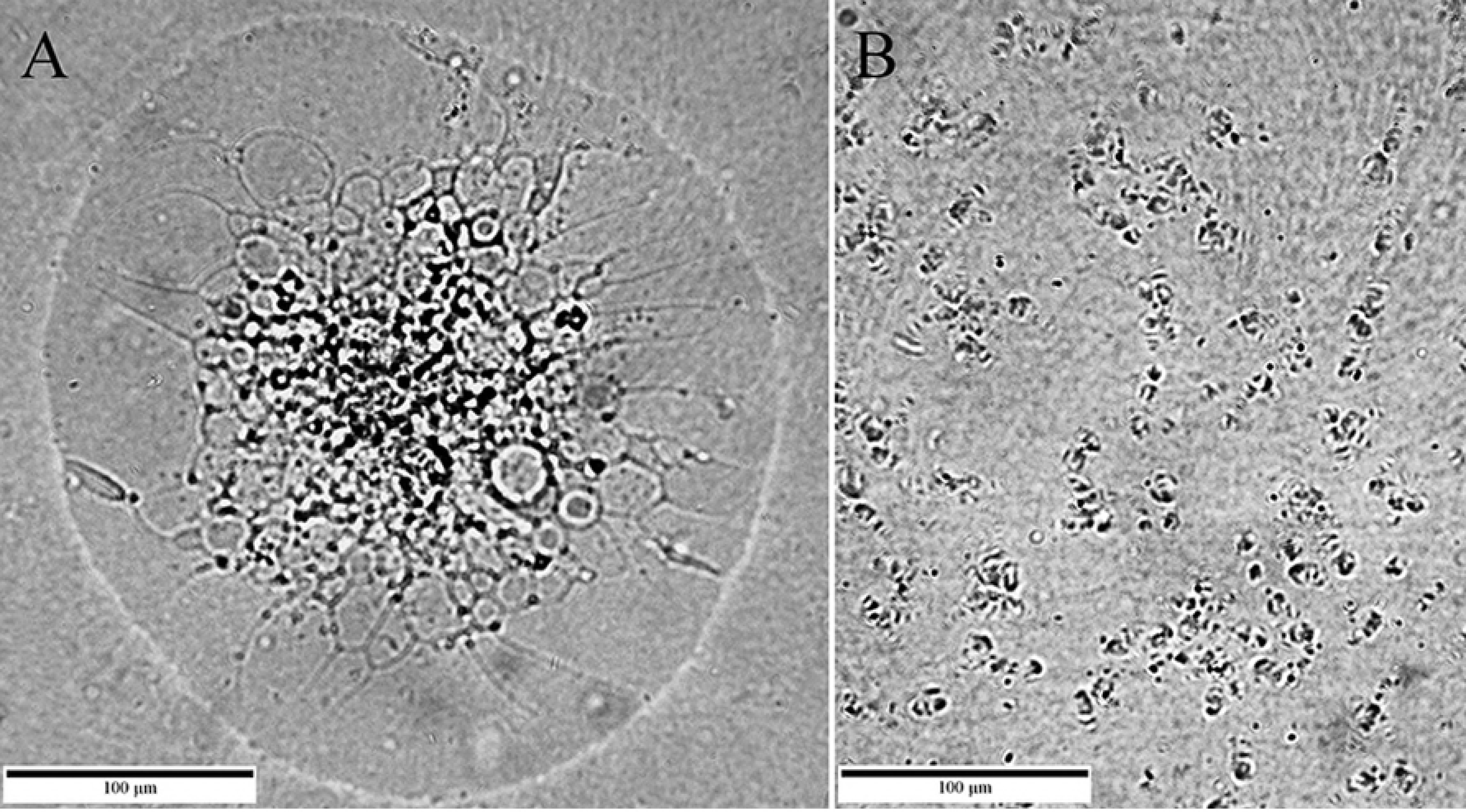
Morphology of M. hominis strain H-34 TC (A) and MCs (B). A. Typical M. hominis colony with the “fried-egg” morphology. B – MCs derived from the M. hominis H-34 culture treated with 50 % hyperimmune serum as described in Materials and Methods and below.

Initial MC cultures obtained from serum samples were re-plated by the “block” method (see Materials and Methods). In some cases, MCs disappeared after 1 or 2 consecutive re-plating steps. Other isolates repeatedly grew and increased in their number with every new re-plating step. These results suggested that we dealt with rather colony multiplication than printing. It is important that consecutive re-plating of MCs have never resulted in appearance of TCs neither in these nor in latter experiments, which are described below.

### Obtaining MCs in vitro

Once MCs were obtained from serum samples, we suggested that interactions with the hyperimmune serum might be a signal to get MCs from the *M. hominis* culture. We treated 10^7^ CFUs of *M. hominis* type strain H-34 grown in the broth for 48 h with dilutions of hyperimmune rabbit antiserum. As it could be predicted, typical *M. hominis* colonies (TCs) were not observed at high concentrations of antiserum (50 and 10 %) but TCs appeared in control and highly diluted antiserum samples on the 3^rd^ day (Fig. 2). In the presence of 0,1 % antiserum, the number of TCs decreased by a factor of 3,7×10^3^up to 1,7 x 10^4^CFU ml^-1^. When 1 % antiserum was used only singular TCs per a plate (50±14 TCs ml^-1^) were observed. In contrast, MCs were observed in all samples including that was treated with 50 % antiserum on the 9th day. The control sample included MCs as well although it was difficult to observe MCs at high concentrations of TCs that seemed to shield MCs from the view. This assay demonstrated that MCs are less or even non-sensitive to hyperimmune antiserum and explained why MCs but not TCs were revealed in clinical serum samples. Similar approach applied to *M. pneumoniae, M. gallicepticum* and *M. fermentans* gave very similar results (Fig. 3).

**Fig. 2.**
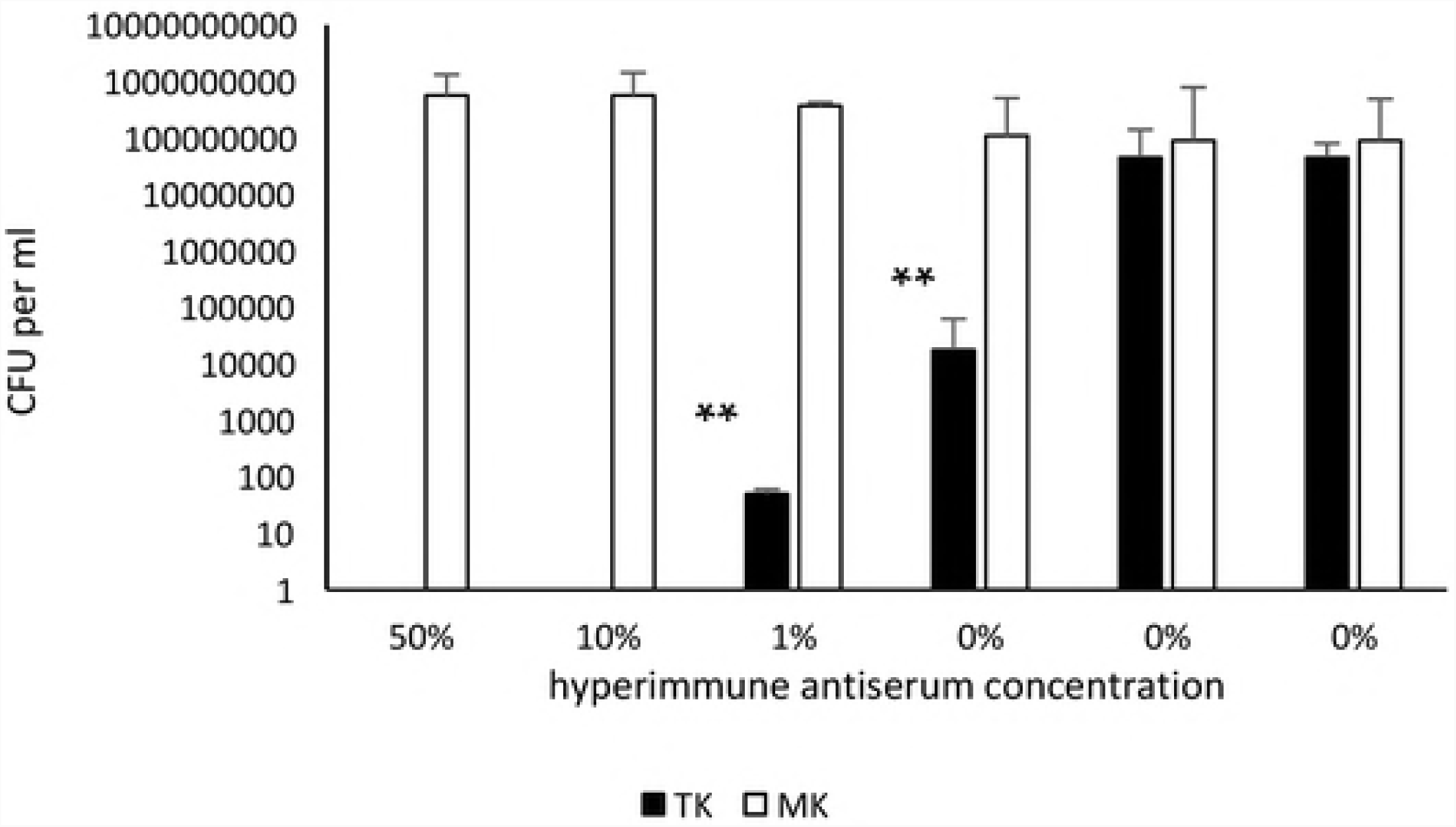
Absolute numbers of TCs and MCs in the *M. hominis* H-34 culture treated with different concentrations of hyperimmune antiserum. The broth *M. hominis* culture containing 10^7^CFU ml^-^(as measured by counting TCs) was incubated with decimal dilutions of hyperimmune antiserum (obtained as described in Materials and Methods) for 24 h. Then decimal dilutions of the culture was plated on the BBL agar. TCs were counted 4 days post plating. MCs were counted 9 days post plating. The data represent mean values ±SD. * p<0,05; ** p<0,01.

**Fig. 3.**
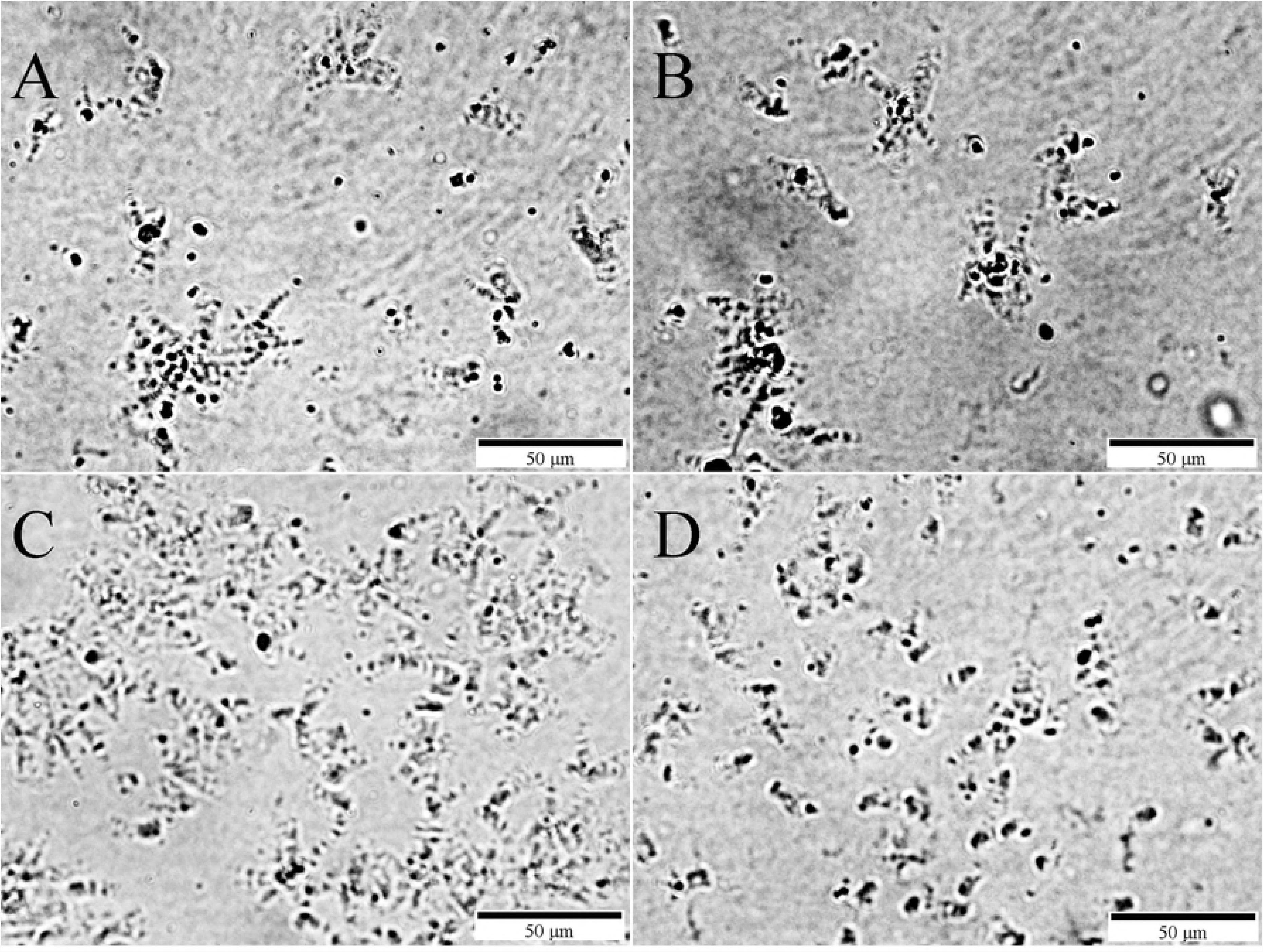
Pure MC culture of different *Mycoplasma* species. MC cultures were obtained with species-specific hyperimmune antiserum treatments as described in Fig. 2. A – *M. fermentans, B - M. gallicepticum*, C - *M. hominis,* and *M. pneumoniae.* Culture were grown for 14 days. Images were taken with magnification 300x

Another method we used to obtain MCs was treatment of *Mycoplasma* spp culture with non-thermal argon plasma. Previously, we observed MC appearance in the *M. hominis* culture treated with argon non-thermal plasma generated by the microwave plasma source MicroPlaSter β developed for the wound therapy [24,25]. We applied non-thermal plasma to treat *M. hominis* and other *Mycoplasma* spp, including *M. pneumonia, M. gallisepticum* and *M. fermentans*. MCs appeared in treated samples of all species. Morphologically, MCs formed by different species were similar. Further, plasma treatment of *Acholeplasma laidlawii* brought about appearance of MCs, too. We suggested that MC formation might be characteristic not only for *M. hominis*, but this phenomenon might be widely spread among *Mollicutes*.

### Specific tests confirmed that MCs are formed by the same species which they were derived from

To prove that MCs belonged to the same species which they were derived from, we applied a number of tests including consecutive re-plating, immunostaining and additional PCR tests. Here results obtained with *M. hominis* H-34 culture and the MC culture described from H-34 with hyperimmune serum treatment are described. MC cultures obtained from clinical samples that gave a positive signal in the PCR test used for clinical sample characterization, demonstrated similar results as H-34 MCs. Moreover, similar results were obtained for all Mycoplasma species tested (data not shown).

We used method of direct fluorescence to stain the MC culture with *M. hominis* specific fluorescent labelled antiserum (see Materials and Methods). The MC culture gave positive signal in the direct fluorescence test (Fig. 4A). The data were confirmed with epifluorescence methods (data not shown). Then we used a camel-derived nanoantibody aMh-FcG2a kindly provided by Dr. Burmistrova. aMh-FcG2a specifically recognizes *M. hominis* membrane ABC transporter substrate binding protein MH3620 of the phosphonate transfer system [19]. Lipid-associated membrane proteins were purified from the *M. hominis* H-34 TC culture and the corresponding MC culture obtained as described above and probed with aMh-FcG2a (Fig. 4B). The MH3620 protein was revealed in both cultures.

**Fig. 4.**
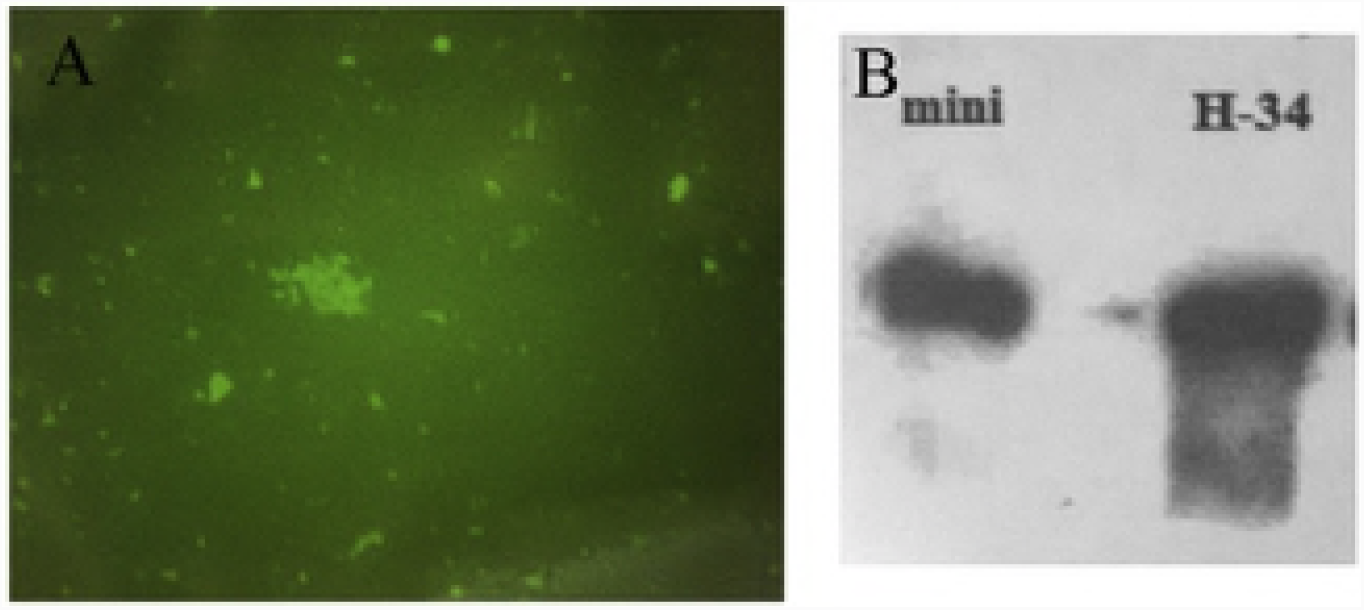
Species-specific tests of MCs derived from the *M. hominis* H-34 culture. A – direct immunofluorescence test of MCs reprinted at the glass; B – western blotting of LAMPs (lipid-associated membrane proteins) isolated from the pure MC culture; LAMPs were labelled with the a camel-derived nanoantibody aMh-FcG2a (Burmistrova et al., 2016).

We determined sequence of 16S rRNA on the DNA isolated from *M. hominis* H-34 TCs and corresponding MCs. Both TCs and MCs had identical 16S rRNA. Then, we applied PCR to reveal 4 housekeeping genes: *alaS, secY, hsdR* and *lysS*. The genes are located on the opposite sides of the *M. hominis* chromosome. Both TCs and MCs gave similar signals in PCR with all 4 primer pairs (data not shown) suggesting that MCs are formed by *M. hominis*.

### Characterization of MC growth and morphology

We followed growth of the pure MC culture obtained from the *M. hominis* type strain H-34 by plasma treatment. MCs were re-plated by the block method. The first appearance of MCs started on the 4^th^ when they strongly resembled agar defects under the microscope with magnification 300x (Fig. 5). On the 7^th^ day MCs were clearly visible as oval or elongated bumps. Since 7^th^up to 12^th^ day colonies increased in their size and took their final morphology. In contrast to round-shaped TCs, MCs had a form resembling a propeller or a spiral that made evident by the 12^th^ day. MCs kept growing after the 12^th^ day although changes in their size or morphology were rather minimal.

**Fig. 5.**
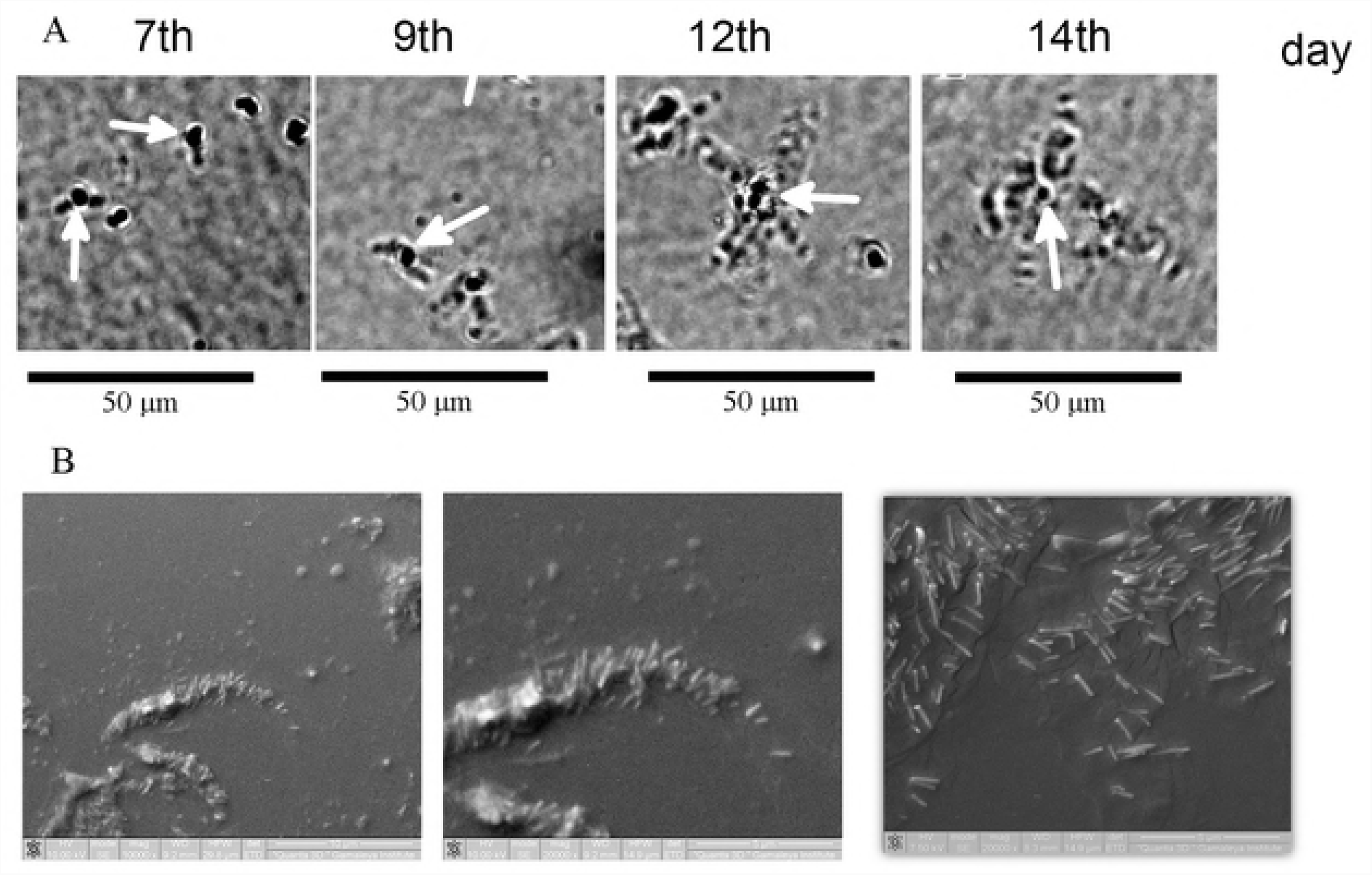
Growth and morphology of MCs. A – *M. hominis* H-34 MCs were plated on the BBL agar and photos were taken with light microscopy (magnification 300x) on 7, 9, 12 and 14 days. Colony development from bulge to propeller-like structure was observed. B – SEM structure of 14 day old MC with magnifications of 10 000x (left) and 20 000x (central and right). Rod-shaped bacterial cells are clearly observed at higher magnification.

The scanning electron microscopy confirmed the propeller-shaped form of MCs (Fig. 5B). Colony centers formed bulges that appeared to be submerged into agar. Under zoom, it is evident that both wings and a center of the propeller are formed by extended rods arranged in parallel and forming spiral forms. The average diameter of rods was about 100 nm, while the length ranged from 200 to 1000 nm with a predominance of 500-600 nm.

### Biochemical characterization

Organisms included in *Mollicutes* are divided into two groups according to their ability to metabolize glucose which are fermenting and non-fermenting mycoplasmas[26,27]. The nonfermenting species hydrolyze arginine. Tests for metabolism of glucose and arginine are the obligatory in mycoplasma characterization. The tests are based on the determination of color change of medium with pH indicator (phenol red)[13]. Particularly, glucose hydrolysis is a typical feature of *M. pneumoniae*. The color of the indicator when M pneumoniae TC culture was added changed from pink to yellow (Fig. 6). In contrast to TCs, the MC colony culture did not change the medium color that suggested inability of MCs to hydrolyze glucose.

**Fig. 6.**
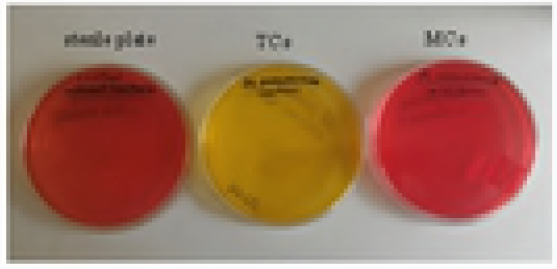
*M. pneumoniae* FH MCs does not hydrolyze glucose. The classic M. pneumoniae culture and corresponding MC culture were plated on the BBL agar supplemented with 1 % glucose and phenol red dye. Changes in the dye color was monitored on 14^th^day for both cultures. Plates from the left to the right: control sterile plate, TCs, MCs.

*M. hominis* is an arginine hydrolyzing species. Arginine hydrolase activity of the TC culture *M. hominis* changed the indicator medium color from yellow to red but no color change was observed for the corresponding MC culture (data not shown). Then we tested *M. hominis* H-34 MC growth in dependence on the presence of arginine. TCs and MCs were subjected multiple re-culturing on the medium supplemented or not with 1% arginine. In the absence of arginine, TCs were re-cultivated up to the 7^th^ passage to disappear at later passages. In contrast, MCs were viable up to 10 passages without noticeable changes in quantity and quality (data not shown). Obtained results suggested that arginine requirements of MCs are lower than requirements of TCs.

### Antibiotic resistance

Comparative resistance to antibiotics was tested for TCs and MCs. Type strains of 4 species including *M. hominis, M. pneumoniae, M. gallisepticum* and *A. laidlawii.* Additionally, the clinical *M. hominis* isolate Izmaylova was included. *M. hominis* and *M. pneumoniae* MC cultures were obtained from the routinely cultivated cultures by hyperimmune serum treatment as described above. *M. gallisepticum* and *A. laidlawii* MCs were obtained by culture treatment with argon gas non-thermal plasma. Initial cultures were used for comparison. TCs were sensitive for the majority of antibiotics tested (Table 2). Only M. hominis strains were resistant to erythromycin. M. hominis was low sensitive to norfloxacin and clarithromycin, M. gallisepticum and A. laidlawii were low sensitive to norfloxacin, gentamycin and clarithromycin, and norfloxacin and doxycycline, respectively. Meanwhile, MCs of all species were resistant to all 9 antibiotics tested.

**Table 1.** MC strains isolated from clinical samples.

|  | Species | source | n |
| --- | --- | --- | --- |
|  | M. hominis n=70 |  |  |
|  |  | serum | 49 |
|  |  | urina | 13 |
|  |  | urinary stones | 5 |
|  |  | Synovial liquid | 3 |
|  | M. pneumoniae, n=2 | Serum | 2 |
|  | M. fermentans, n=2 | Synovial fluid | 2 |
|  | Mycoplasma spp, n=5 | Serum | 3 |
|  |  | Synovial fluid | 2 |

**Table 2.** Antibiotic resistance of TCs and MCs

| Group<sup>a</sup> | Antibiotic | Dose<br>(µg) | *M. hominis*<br>H-34 |  | *M. hominis*<br>Izmaylova |  | *M. pneumoniae* |  | *M. gallisepticum* |  | *A. laidlawii* |  |
| --- | --- | --- | --- | --- | --- | --- | --- | --- | --- | --- | --- | --- |
|  |  |  | TC | MC | TC | MC | TC | MC | TC | MC | TC | MC |
| Ag | Gentamycin | 10 | +<sup>b</sup> | -<sup>d</sup> | + | - | + | - | ± | - | + | - |
| Flu | Norfloxacin | 10 | ±<sup>c</sup> | - | ± | - | + | - | + | - | ± | - |
| Tet | Doxycycline | 10 | + | - | + | - | + | - | + | - | ± | - |
| Macr | Clarithromycin | 15 | ± | - | ± | - | + | - | ± | - | + | - |
|  | Roxytromycin | 30 | + | - | + | - | + | - | + | - | + | - |
|  | Clyndamycin | 2 | + | - | + | - | + | - | + | - | + | - |
|  | Azythromycin | 15 | + | - | + | - | + | - | + | - | + | - |
|  | Erythromycin | 15 | - | - | - | - | + | - | + | - | + | - |
| Lin | Linkomycin | 15 | + | - | + | - | + | - | + | - | + | - |

<sup>a</sup> Antibiotic groups: Ag -aminoglycoside group, Flu -fluoroquinolone group, Tet - tetracycline group, Macr – macrolide group, Lin – lincosamide group.
<sup>b</sup> sensitive
<sup>c</sup> intermediate (see in Materials and methods)
<sup>d</sup> resistant

Moreover, when antibiotic supplemented broth cultures were seeded with the routinely cultivated cultures, TCs were eliminated and pure MCs were grown upon agar culture plating. Therefore, besides treatment with the hyperimmune serum and non-thermal argon plasma, antibiotic treatment might help to isolate a pure MC culture.

### Typical colonies carry MC forming units

Obtained results demonstrated that MCs might be obtained from the TC culture by treatment with the hyperimmune serum, non-thermal atmospheric plasm or antibiotics. Our question was whether it was antibacterial treatments that determined MC development. Alternatively, MCs forming units could persist within a TC to survive treatments, which are non-permissive for TC forming units. To address this question, we analyzed an isolated TC, which was not treated in any way. We used a method of colony growth in semiliquid agar to get an isolated TC of *M. hominis* H-34. This method allowed to get an intact *M. hominis* colony while otherwise the part of colony immersed into agar would be lost. The isolated colony was removed from semiliquid agar, suspended in 1 ml of the broth and plated on standard agar. Plates were incubated for 12 days. All formed colonies were counted on the 2^nd^, 9^th^ and 12^th^ days (Fig. 7A). TCs were appeared as soon as in 24-48 h, and their number did not increase starting from the 48 h. After 120 h (5^th^ day), the number of TC gradually declined as a part of TCs underwent a lysis. MCs got visible (with the magnification 300x) on the 4^h^ day, and reached maximal count on the 12^th^ day (with the magnification 120x). The relative number of MCs to TCs on the 12^th^ day was 4,8±2,1. Obtained results demonstrated that a pure MC culture might be obtained from a TC without any treatment.

**Fig. 7.**
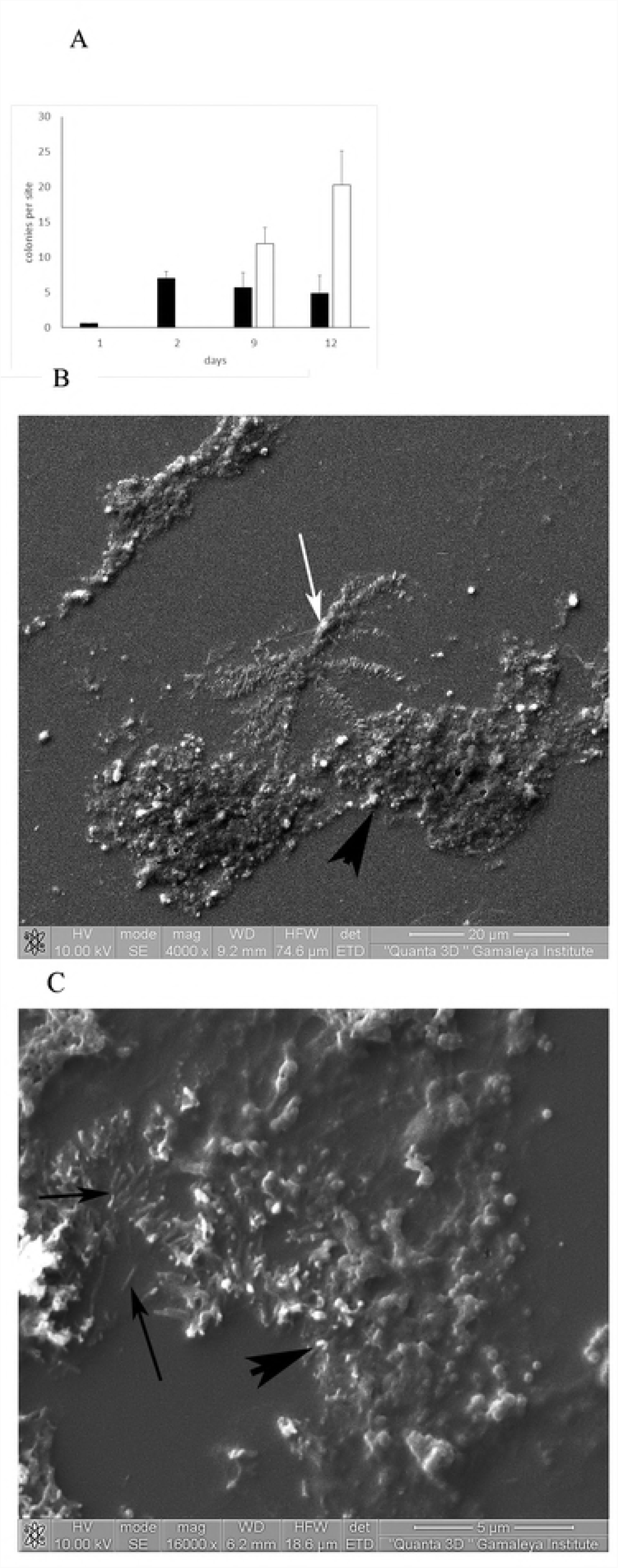
Identification of MCs in the routinely M. hominis H-34 culture. A – absolute numbers of TCs and MCs plated from the isolated colony grown in the semiliquid agar. B – MC (white arrow) and fried-egg type TC (black arrowhead) located in the close vicinity observed with SEM (magnification 4000x). C – rod-like cells (arrows) typical for MCs were observed along with typical cell forms (arrowhead) within a typical colony (TC).

Then we thoroughly studied the type strain H-34 *M. hominis* culture with SEM. The culture was routinely re-plated many times since it was obtained from Dr. Lemke in 1967 and used as a control in all studies. Both TCs and MCs were observed within the culture with SEM as well as with the light microscopy (Fig. 7B). TCs when observed under the higher resolution included predominantly round-shaped variable size cells (Fig. 7C). Meanwhile, in some cases, groups of elongated roads similar to MC cells were observed within TC (Fig. 7C). Taken together, these results suggested that MC forming units might be different from TC forming units and both were present within a typical colony. In other words, our results suggested that TC is primarily heterogenous.

### Occurrence of MCs in clinical samples

Our last task was to establish how wide the phenomenon of MCs is spread. As stated above, firstly we revealed MCs in sera of patients that were tested on *M. hominis*. Totally, 75 samples PCR-positive samples obtained from patients with inflammatory urogenital tract diseases were studied. MCs were isolated from 37 (49,2 %) of them. Viability of MCs was confirmed by re-plating. The majority of samples were re-plated 2 or 3 times after that the culture lost its viability. Meanwhile, some cultures were re-plated repeatedly more than 10 times and maintained their viability for at least 3 years. MCs have not changed their morphology in the course of re-plating for 3 years, and TCs have never formed.

To address MC spreading in other specimen, we looked for MCs in urina and urinary stones of patients with nephrolithiasis. PCR-positive signals suggesting *Mycoplasma* spp persistence was found in samples of 25 of 31 patients (Table 3). Plating of PCR-positive samples revealed MCs in samples obtained from 17 of 25 patients with PCR-positive samples. Best matching between PCR results and MC isolation was found for samples of urina: MCs were revealed in 8 of 9 PCR-positve samples (88,8 %). The worst correlation was found for samples of crushed urinary stones: MCs were grown from only 5 of 21 (22,8 %) PCR-positive samples. There were no TCs grown from the studied samples. Absence of TCs could be due to extensive antibiotic treatments of patients before and after urinary stone fragmentation.

**Table 3.** Occurrence of MCs in samples obtained from patients with urolithiasis

| Sampling | *Mycoplasma*<br>spp PCR-positive (%) | MCs in<br>PCR-positive<br>samples<br>(%) | TCs in<br>PCR-positive<br>samples |
| --- | --- | --- | --- |
| Patients, 31<br>Including | **25** (100 %) | **17** (68 %) | **0** |
| Urina, 31 | **9** (100 %) | **8** (88,8 %) | **0** |
| Urina from renal hilus,<br>31 | **14** (100 %) | **5** (35,7 %) | **0** |
| Urinary stones, 31 | **21** (100 %) | **5** (22,8 %) | **0** |
| Serum, 31 | **8** (100 %) | **5** (62,5 %) | **0** |

Next, we addressed dissemination of MC formed by *Mycoplasma* spp among patients with other pathologies. Particularly, we performed bacteriological analysis of synovial fluid and serum samples obtained from patients with rheumatoid arthritis and asthma, respectively. Both, samples positive and negative in PCR on *Mycoplasma* spp. were plated. 10 of 25 synovial fluid samples gave a positive signal in PCR (Table 4). MCs were isolated from 7 of these 10 samples. *Mycoplasma* spp were not found in PCR-negative samples. No one of samples included TCs. Further, we studied 74 samples from patients with asthma. *Mycoplasma* spp specific positive PCR-results were obtained for 31 samples (42 %). MCs were plated from 10 of these samples.

**Table 4.** Occurrence of MCs in samples obtained from patients with rheumatoid arthritis and asthma

| Samples<br>(pathology, N) | *Mycoplasma*<br>spp PCR-positive | MCs in |  | TCs in |  |
| --- | --- | --- | --- | --- | --- |
|  |  | PCR-positive<br>samples | PCR-negative<br>samples | PCR-positive<br>samples | PCR-negative<br>samples |
| synovial fluid<br>(rheumatoid<br>arthritis, 25) | 10 | 7 | 0 | 0 | 0 |
| Serum<br>(asthma/bronchitis;<br>74) | 31 | 10 | 0 | 0 | 0 |

Taken together, the total number of MC strains isolated from clinical specimens was 79, including 70 *M. hominis*, 2 *M. fermentans*, 2 *M. pneumoniae* and 5 *Mycoplasma* spp strains. Prevalence *of M. hominis* might be related to both wide spread of this pathogen in human population and less insistence of this microorganism to cultivation conditions.

## Discussion

Results presented here demonstrated a novel previously undescribed type of Mycoplasma colonies that were designated as microcolonies (MCs) as distinct from typical colonies (TCs). MCs are formed by cells that are morphologically distinctive from normal Mycoplasma cells in SEM. MCs were observed within replated cultures of type strains of at least five Mycoplasma species. and MC forming cells were observed within typical colonies of these species. Careful consideration of Mycoplasma monolay microphotos in the basic textbooks and articles could reveal MCs that grow between TCs. Thus, one can speak about primary heterogeneity of *Mycoplasma* population that includes typical cells and MC forming cells.

Pure MC cultures were obtained in vitro under stress conditions such as specific antiserum, antibiotic application or treatment with non-thermal gas plasma applied to TC cultures. Moreover, pure MC cultures were obtained from clinical samples of blood, plasma, urina etc. which were Mycoplasma positive in PCR. Noticeably, pure MC cultures were obtained from patients with different pathological often chronic diseases including urogenital tract diseases, bronchial asthma, rheumatoid arthritis. MCs were as well isolated from the blood of different species of monkeys, cryoconserved bull sperm and cell cultures (data not shown). Taken together, these data suggest wide spreading of Mycoplasma MCs.

The major differences between TCs and MCs included (i) colony morphology and size: TCs have a “fired-egg” look and consist of rounded cells of different sizes and single rod-shaped cells; MCs have a propeller like form formed by rows of paralleled rod-shaped cells. TC diameters ranged from 50 to 800 µm while MC diameters were within 7,5-9,5 µm; (ii) growth rates: TCs grew for 48-96 h for all Mycoplasma species tested while MCs got visible under the magnification 120x on the 7-9 and up to 12th day post plating; (iii) MCs hydrolyzed neither glucose nor arginine or at least hydrolysis was not observed with standard tests; (iv) MCs were resistant to stresses that eliminated TCs; (v) MCs were resistance to 9 antibiotics belonging to different classes.

It seems that small colony sizes and very slow growth rates have hided MCs from observations up to date. Then what are MCs? To make an unambiguous answer, the proteomic analysis and genome comparison should be performed that allows detection of genes and proteins responsible for their specific morphotype, physiology and antibiotic resistance. Preliminary data suggest that TC and MC proteomes are substantially different. The proteomic analysis of *M. hominis* TCs revealed 346 proteins versus 291 proteins in the MC culture obtained with immune treatment as described here [28]. Still, particular molecular mechanisms responsible for existence of MCs have not been established yet. In the absence of this information, one can make a few conclusions based on the presented data.

It is clear that MCs are not artifacts known as “pseudocolonies” that were described in early works [13]. These pseudocolonies that are very similar to Mycoplasma TCs have a solid center and filamentous periphery similar to TCs when observed with light microscopy. However, more detailed analysis revealed that these were eukaryotic cells, or aggregates formed by detergents and/or phosphoric or calcium salts of fatty acids. The phenomenon of MCs is different from these artifacts.

Bacterial cells that form MCs are not nanobacteria. Nanobacteria were described in the beginning of XXI century as bacteria with the cell size of 100-150 nm that can not be seen with light microscopy [29]. Data obtained with electron microscopy applied to study nanobacteria suggested that the observed nanocells were rather crystals of apatite or calcium salts [30]. Up to date, nanobacteria were not cultivalable that allowed suggestion about nanobacteria as a VBNC (viable but not cultivable) form of Mycoplasmas. MC forming cells are not nanobacteria as although their diameter ranged 100-120 nm, their length reached 600 nm, they grow on agar and could be replated many times.

MC forming cells cannot be described as persistors, which are dormant transiently non-dividing cells present in populations of majority of bacterial species in proportion of up to 1-3 % [31–33]. MC forming cells divided and they did restore TC phenotype.

According to their relative sizes, MCs could have been assigned to so-called “small colony variants” (SCVs)[34,35]. SCVs were described in many bacterial species including *Salmonella typhi, Escherichia coli, Pseudomonas aeruginosa, Staphylococcus* spp. SCVs were suggested to be linked to chronic, recurrent, and antibiotic-resistant infections. Molecular mechanisms underlying SCV formation were well defined for *S. aureus* where mutations affecting electron transport, small regulatory RNAs, tymidyl, hellicase and L6 ribosomal protein synthesis were shown to be associated with the SCV phenotype[34,36,37]. In other words, one can speak about genetic divergence in bacterial populations that results in survival of a part of the population under selective pressure that maybe exerted by immune system, antibiotic treatment etc. The Mycoplasma colony size was shown to be dependent on cultivation medium, a loading dose, development of the nearest colonies [12]. However these variations have a transient character, and colonies restore their size when conditions improve. The phenomenon of the MCs is different, because the MC phenotype is stable and hereditary. Therefore, we suppose that appearance of MCs might be a result of genetic divergence as it was shown for some other bacterial SCVs. Meanwhile, MCs formed by Mycoplasmas seem to be more stable than SCVs formed by other bacterial species. Generally, SCVs have high rates of reversion to the normal phenotype under in vitro conditions that might be strongly selected for due to the increased growth rate of normal size colonies on laboratory media [34–38]. Reversion rates is an important characteristic of SCVs. Precise mechanisms of reversion have not been established for all mutations although some studies demonstrated that compensating mutations at the initially identified mutation site are responsible for the reversion phenotype that does not exclude alternative mutation sites [36]. For *A. laidlawii* and *M. gallisepticum*, mutations appeared under antibiotic pressure or starvation result in morphologic changes and a shift in proteomic patterns [39,40]. Still, the phenotype was not stable and reverted to the normal one when the antibiotic pressure was stopped. Meanwhile, Mycoplasma MCs described here were exceptionally stable. Regular replating for up to 3 years, cell culture and laboratory animal infection (data not shown) did not result in reversion of MCs to TCs. These data suggested that appearance of MCs might be a result of genetic divergence due to multiple mutations that prevents reversion. This suggestion is in line with relatively high mutation rates that differ Mycoplasmas from other bacterial species. Certain analogy might be given with tumor cells that appeared due to multiple mutations in somatic cells for never to revert to the initial genotype.

Obtained results suggested that *Mycoplasma* population is heterogeneous in absence of selective pressure. Particularly, this suggestion is supported by the following observations: repeatedly obtained typical colonies grown in semiliquid agar, resuspended, diluted and replated gave both TCs and MCs every time; SEM demonstrated presence of rod-shaped cells characteristic for MCs within a TC or in close vicinity to TCs. These data suggested that external factors such as hyperimmune antiserum, antibiotics or non-thermal gas plasma rather killed TC cells saving MC cells alive than stimulated MC development from TCs. Still, a stimulatory potential of the factors listed can not be excluded totally. Then the question rises why MCs do not eliminate from the population as they seemed to have slower growth rates. The one answer is that we still do not exactly know how different division rates are for cells forming TCs and MCs. The slow appearance of MCs for the observation might not be fully due to slow division rates but rather due to tight packing of cells while division rates might be comparable for TC and MC cells. Anyway, presence of MC cells within Mycoplasma population, their viability in conditions when TCs were not observed suggested that MC formation gives some evolutionary advantage to Mycoplasma species.

One of the most intriguing results obtained was resistance of MCs to antibiotics with different mechanisms of antibacterial activity. Mutations that provide resistance in Mycoplasmas to each of the studied antibiotics are established. Macrolide resistance is caused by point mutations of the macrolide-binding site located in the 23S rRNA genes or in the genes encoding ribosomal proteins L4 and L22 [41,42]. Point mutation(s) in the 16S rRNA genes was described for tetracycline resistant mutants of *M. pneumoniae* [43] and *M. bovis* [44]. The only mechanism of fluoroquinolone resistance described in Mycoplasmas is alterations within the quinolone resistance-determining regions (QRDRs) of DNA gyrase subunits GyrA and GyrB and/or topoisomerase IV subunits ParC and ParE [41]. Meanwhile, MCs were resistant for all antibiotics tested that suggested a general mechanism of resistance. This mechanism might be due to changes in MC membrane composition or absence of certain transporter proteins.

The key question for further studies of Mycoplasma MCs is their role in human infectious pathology. The important role in bacterial persistence in the human organism was suggested for S. aureus SCVs [34,45]. This role is particularly due to an ability of SVC bacteria to survive unfavorable conditions and is supported by further SCV reversion into wild type forms. According our observations, MCs do not revert in TCs and therefore the role in pathology should be studied for MCs themselves and this is a challenge for the future. Meanwhile, obtained data are important for understanding features important for optimization of modern systems for Mycoplasma spp diagnostics. Antibiotic treatment of patients that suggested to be Mycoplasma spp infected on the basement of the PCR analysis only might be wasteful as MC forming bacteria that can give PCR-positive results are fully antibiotic resistant. On the other hand, existing biochemistry-based test-systems for Mycoplasma identification in clinical samples or tissue cultures are useless for detection of MCs as they do not hydrolyze neither glucose nor arginine.

Taken together, obtained results described a novel form of pathogenic *Mycoplasmas*. This form is characterized by specific cell and colony morphology, changes in key biochemical markers, unspecific resistance to antibiotic treatments and other stresses including immune response, and a wide spread in clinical samples. Elucidation of the role that this form might play in infectious disease might be an important step in more deep understanding of mechanisms used by bacterial pathogens in development of persistent infections.

## Acknowledgements

Authors are thankful to Dr. K.H. Lemcke (Listar Instute of Preventive Medicine, London, UK), Dr R.M. Chanock (National Institue of Health, Beathesda, USA), Dr E.A. Freundt (Institute of Medical Microbiology, University of Aarhus,Denmark), Dr. K.P. Koller (Germany) who provided Mycoplasma type strains to Prof. G.Ya. Kagan, the former leader of our lab. Authors thank Dr. Burmistrova for the M. hominis specific aMh-FcG2a nanoantibody. The non-thermal argon plasma MicroPlaSter source was generously provided by Prof. G.E. Morfill (MPE, Germany) and Adtec Plasma Technologies Ltd (Japan).

## Materials and Methods

### Ethical statement

All experiments utilizing rabbits were performed in compliance with guidelines established by the Animal Welfare Act for housing and care of laboratory animals, with the Russian Federation National Standard (GOST R52379-2005), directives of Ministry of Health of Russian Federation (No753n from 26.08.2010, No774H from 31.08.2010), and with the approval of the Biomedical Ethics Committee of Gamaleya Research Center of Epidemiology and Microbiology (No9a from 24.01.2017). Rabbits were fed standard rabbit chow daily. Appropriate measures were utilized to reduce potential distress, pain and discomfort. All animals received environmental enrichment.

Acquisition of the patient samples used in this study was approved by the Gamaleya National Research Center of Epidemiology and Microbiology (Moscow) and by the “Finist” Ltd (Moscow), S.P. Botkin hospital (Moscow), hospital of the 1^st^ Moscow Medical University named after I.M. Sechenov, and all patients or their representatives gave written, informed consent.

### Bacterial strains and media

The following *Mycoplasma* type strains were used in the study: the *M. hominis* strain H-34 kindly provided by Dr. K.H. Lemcke (Listar Instute of Preventive Medicine, London, UK); the *M. pneumonia* FH strain, kindly provided by Dr R.M. Chanock (National Institue of Health, Beathesda, USA), the *M. gallisepticum* strain S6 and the *M. fermentans* strain Pg18, were kindly provided by Dr E.A. Freundt (Institute of Medical Microbiology, University of Aarhus,Denmark). *Acholeplasma laidlawii* were kindly provided by Dr. K.P. Koller (Germany). Isolates obtained in this study are listed in the Table 1.

*Mycoplasma* spp were cultivated on BBL™ Mycoplasma agar (PPLO Agar base, BD, USA) and in the Difco™ PPLO Broth (BD, USA) supplemented with 15 % horse serum (Gibco, ThermoFisher Scientific, USA) and 1 % L-arginine (for *M. hominis*) or 1% glucose for other species. Penicillin was added up to 100 µg ml^-1^ to avoid contamination.

### Growth conditions

Colonies that appeared on BBL™ Mycoplasma agar plates were observed with the light microscope with objectives 10X, 25X, 40X (LETZLAR, Germany). TCs were counted 96 h post plating. MCs were counted when they got clearly visible, usually on the 7^th^-9^th^ day.

To re-plate MCs we applied a “block method”. Shortly, a cube of agar with a side of 1 cm was cut off from the plate with MCs. To evenly distribute bacteria, the cube was applied its upside down to a fresh plate with a light circular motion.

### Mycoplasma spp isolation from clinical samples

All clinical samples were obtained and studied in the frame of performance of laboratory tests prescribed by a hospital attending physician, and further prolonged incubation of the samples did not require a special permission. The following clinical samples were used. 75 serum samples were obtained from patients with inflammatory urogenital tract diseases (urethritis, prostatitis, cervicitis) surveyed on *M. hominis* at “Finist” Ltd (Moscow, Russia). 124 samples were obtained from patients with nephrolithiasis (n=31) subjected to nephrolitholapaxia at S.P. Botkin hospital (Moscow, Russia). Primary urine (Urine1, n=31), urine obtained by catheterization from renal pelvis (Urine2, n=31), serum (n=31) and lithotripsyc stones (n=31) were used. 25 samples of synovial join fluid were obtained from children 2 to 16 years old with diagnosis of arthritis with aggressive joint syndrome that were treated at hospital of the 1^st^ Moscow Medical University named after I.M. Sechenov. 74 samples of serum from patients with chronic diseases of respiratory tract (chronic bronchitis, asthmatic bronchitis, asthma) were obtained at the same hospital.

According to a standard procedure, all samples were firstly studied with PCR with the universal “MYCOM” test-system (InterLabService Ltd, Moscow, Russia). Positive samples were further studied. To differentiate Mycoplasma spp, “Amplisens-Mycoplasma hominis E-ph”, “Amplisens-Mycoplasma gentialium”, Amplisens-Mycoplasma pneumoniae” were purchased from the same manufacturer. To differentiate *M. fermentas*, biochemical tests were used. Then all PCR-positive samples were studied in a bacteriological assay by using BBL™Mycoplasma agar plates. Microscopic plate evaluation was performed as described above.

### Arginine- and glucose-hydrolysis testing

Arginine and glucose-hydrolysis was detected on the BBL agar supplemented with 1% L-arginine (Dia-M, Russia) or 1 % glucose, respectively, and 0,03 % phenol red (Sigma-Aldrich, USA). Mycoplasma were plated by a “cube” method as described above. Plates were incubated for two weeks at 37°C.

### Antibiotic resistance

The disc diffusion antibiotic susceptibility test was performed with discs supplied by “Medica+ Ltd” (Sankt-Petersburg, Russia). Results were presented as “+” (sensitive) – an inhibition zone without growth; “-“(resistant) – full growth; “±” (intermediate) – single colonies were observed within the inhibition zone at a distance of 0,5-1 cm from the disc.

### Protein purification, SDS-PAGE and Western-blotting

*M. hominis* lipid-associated membrane proteins (LAMPs) were purified mainly as described in [19]. Shortly, TC and MC cultures were washed away from agar plates (5 and 30 plates were used, respectively), Bacteria were precipitated by centrifugation at 10000 g for 20 minutes. Cells were resuspended in the TE-TX114 buffer (50 mMTrisHCl; 150 mM NaCl, ImMEDTA, ph 8,0, 2% Triton X-114) and incubated with constant shaking at 4°C for 2 h. The lysates were centrifuged at 5000 g for 5 minutes at 4°C to remove debris. Supernatant was heated up to 37°C to stimulate formation of lipid micelles. Micelles were separated from the water phase by centrifugation at 5000 g for 5 minutes at 37°C. Then micelles were suspended in TE buffer (50 mMTrisHCl; 150 mMNaCl) at 4°C and repeated a stage of separation of micelles and the water phase twice. Obtained micelles were stored at −20°C. LAMPS were extracted from micelles by addition of an equal volume of methanol followed by centrifugation at 10000 g for 20 minutes. The protein pellet was washed with ethanol for three times and resuspended in PBS. Obtained LAMPs were separated on 12 % SDS-PAGE and transferred on the nitrocellulose membrane. Immunoblotting was performed with the nanoantibody aMh-FcG2a specific to ABC transporter substrate binding protein of the phosphonate transfer system MH3620 [19].

### Development of hyperimmune serum

The hyperimmune serum was developed by rabbit immunization with mycoplasma membrane fractions that were obtained as described in [20]. Shortly, type Mycoplasma strains were grown in the broth for 3 days at 37 °C. Bacteria were precipitated at 9 000 rpm for 45 minutes, resuspended in PBS and disrupted with the Techpan ultrasonic disintergrater for 20 minutes (10 2-minutes cycles with 2-minutes intervals). Membrane fraction was obtained by centrifugation at 40 000 g for 30 minutes. Immunization of 2,5-3 kg rabbits was performed according to the standard scheme [21].

### Direct immunofluorescence and epiimmunofluorescence tests

These tests were performed as described in [22,23] with “MycoHomoFluoScreen” test-system (MedGamal Ltd, Moscow, Russia). Pure MC cultures were studied in comparison with TC cultures. For direct immunofluorescence assay, reprints of mycoplasma colonies grown on agar were obtained on the glass, fixed by 96^°^ ethanol, further glasses were treated with “MycoHomoFluoScreen” test-system as described by manufacturer. For epiimmunofluorescence, colonies grown on agar plates were directly treated with the same test-system and observed with a microscope equipped with an incident illumination module.

### DNA purification and PCR

DNA was obtained from clinical samples with the “DNA-sorbV” kit. For laboratory purposes, bacteria were washed from plates with PBS and lysed by boiling for 5 minutes. Primers were developed against housekeeping genes *alaS, secY, hsdR* and *lysS* (Table 1S). PCR was performed in the Tertcik amplificator (DNA-Technologies, Russia) at the annealing temperature of 55 °C for all primers pairs. Commercial test systems were used according manufacturer’s instructions.

### Mycoplasma treatment with the hyperimmune serum

The broth cultures were used. Full rabbit serum or its corresponding dilutions were added to the bacterial suspension in the ratio 1:1 to get final concentrations described in the text. The mixture was incubated at 37°C for 24 h. Than decimal dilutions were plated on BBL™Mycoplasma agar. TCs were counted 4 days post plating. MCs were counted 12 days post plating.

### Mycoplasma spp treatment with non-thermal plasma

The MicroPlaSter argon microwave plasma source was used [24]. Bacterial treatment was performed as previously described [25]. Briefly, 100 µl of the broth culture (approximately 10^7^ CFU ml^-1^ as determined by counting of TCs 48 h post plating) was plated on the on BBL™. Immediately after plating the culture was treated with argon microwave plasma from the distance of 2 cm for 5 minutes. Then the plates were incubated until MC appearance.

### Scanning electron microscopy (SEM)

Material for SEM was prepared as reprints of colonies on coverslips glass that were fixed by 96^0^ ethanol or Mycoplasma agar-grown colonies that were fixed with 10% formalin vapor for 24h. The samples were coated with 5 nm layer of gold and studied with the dual-bean microscope Quanta 200 3D (FEI Company USA) equipped with the module for spraying SPI-MODULE Sputler Coater (SPI).

### Statistics

All experiments were repeated at least three times. MC re-plating experiments were conducted up to at least 10 consecutive steps with all strains listed in the Bacterial strains section including clinical isolates (n=70). The following MC cultures were maintained by repeated re-plating steps for 3 years: M. hominis H-34 derived MC culture;

The quantitative graphs describing microscopic data are shown as mean values ± SD that were obtained from at least three independent experiments made in 10 vision fields. The mean values and SD were calculated with Excel software (Microsoft Office 2010). The paired ttest included in the same software was used for assessment of statistical significance. Representative light microscopy and SEM photos were chosen from a bulk of similar photos that included more than 50 photos each.

## Author contributions

I.V. R. contributed to conception and design of the study, data acquisition and analysis and manuscript draft writing, G.A.L., O.I.B., A.Ya.M., S.G.A., L.G.G. and E.V.S. contributed to data acquisition and analysis, S.A.E. contributed to data analysis and manuscript draft writing, V.G.Z. and G.G.M. contributed to data analysis, all authors approved manuscript.

## Bibliography

1. Razin S, Hayflick L. Highlights of mycoplasma research-An historical perspective. Biologicals. 2010. doi:10.1016/j.biologicals.2009.11.008

2. Razin S, Yogev D, Naot Y. Molecular biology and pathogenicity of mycoplasmas. Microbiol Mol Biol Rev. 1998; doi:1092-2172/98

3. Hayflick L, Chanock RM. Mycoplasma Species of Man. Bacteriol Rev. 1965;

4. Narita M. Pathogenesis of extrapulmonary manifestations of Mycoplasma pneumoniae infection with special reference to pneumonia. Journal of Infection and Chemotherapy. 2010. doi:10.1007/s10156-010-0044-x

5. Daley GM, Russell DB, Tabrizi SN, McBride J. Mycoplasma genitalium: A review. International Journal of STD and AIDS. 2014. doi:10.1177/0956462413515196

6. Othman N, Isaacs D, Daley AJ, Kesson AM. Mycoplasma pneumoniae infection in a clinical setting. Pediatr Int. 2008; doi:10.1111/j.1442-200X.2008.02644.x

7. Murtha AP, Edwards JM. The role of Mycoplasma and Ureaplasma in adverse pregnancy outcomes. Obstetrics and Gynecology Clinics of North America. 2014. doi:10.1016/j.ogc.2014.08.010

8. Rakovskaia I V, Gorina LG, Balabanov DN, Levina GA, Barkhatova OI, Goncharova SA, et al. [Generalized mycoplasma infection in patients and carriers]. Zh Mikrobiol Epidemiol Immunobiol. 2013;

9. Watanabe H, Uruma T, Nakamura H, Aoshiba K. The role of Mycoplasma pneumoniae infection in the initial onset and exacerbations of asthma. Allergy and Asthma Proceedings. 2014. doi:10.2500/aap.2014.35.3742

10. Yano T, Ichikawa Y, Komatu S, Arai S, Oizumi K. Association of Mycoplasma pneumoniae antigen with initial onset of bronchial asthma. Am J Respir Crit Care Med. 1994; doi:10.1164/ajrccm.149.5.8173777

11. Waites KB, Xiao L, Liu Y, Balish MF, Atkinson TP. Mycoplasma pneumoniae from the respiratory tract and beyond. Clinical Microbiology Reviews. 2017. doi:10.1128/CMR.00114-16

12. Bredt W. Celular morphology of newly isolated Mycoplasma hominis strains. J Bacteriol. 1971;

13. Razin S. Methods of Mycoplasma. In: Razin S, Tully J, editors. New York: Academic Press; 1983. pp. 83–86.

14. Klieneberger-Nobel E. Mycoplasma, a brief historical review. Ann N Y Acad Sci. 1967;143: 713–718.

15. Dybvig K, Simecka JW, Watson HL, Cassell GH. High-frequency variation in Mycoplasma pulmonis colony size. J Bacteriol. 1989; doi:10.1128/jb.171.9.5165-5168.1989

16. Kammer GM, Pollack JD, Klainer AS. Scanning-beam electron microscopy of Mycoplasma pneumoniae. J Bacteriol. 1970;

17. Boatman E. Morphology and Ultrastucture of Mycoplasmatales. In: Barrile M, Razin S, editors. The Mycoplasmas V 1, Cell Biology. New York: Academic Press; 1979. pp. 63–102.

18. Quinlan DC, Maniloff J. Deoxyribonucleic acid synthesis in synchronously growing Mycoplasma gallisepticum. J Bacteriol. 1973;

19. Burmistrova DA, Tillib S V., Shcheblyakov D V., Dolzhikova I V., Shcherbinin DN, Zubkova O V., et al. Genetic passive immunization with adenoviral vector expressing chimeric nanobody-Fc molecules as therapy for genital infection caused by mycoplasma hominis. PLoS One. 2016; doi:10.1371/journal.pone.0150958

20. Razin S, Rottem S. Techniques for the manipulation of Mycoplasma membranes. In: Madd A, editor. Biochemical analysis of membranes. London: Academic Press; 1976.

21. Senterfit L. Preparation of antigens and antisera. Methods in Mycoplasmology, V 2. New York: Academic Press; 1983. pp. 401–404.

22. Del Guidice R, Barile M. Immunofluorescent procedures for mycoplasma identification. Dev Biol Stand. 1974;23: 134–137.

23. Bradbury JM, Oriel CA, Jordan FTW. Simple method for immunofluorescent identification of mycoplasma colonies. J Clin Microbiol. 1976;

24. Shimizu T, Steffes B, Pompl R, Jamitzky F, Bunk W, Ramrath K, et al. Characterization of microwave plasma torch for decontamination. Plasma Process Polym. 2008; doi:10.1002/ppap.200800021

25. Ermolaeva SA, Rakovskaya I V., Miller GG, Sysolyatina E V., Mukhachev AY, Vasiliev MM, et al. Nonthermal plasma affects viability and morphology of Mycoplasma hominis and Acholeplasma laidlawii. J Appl Microbiol. 2014; doi:10.1111/jam.12445

26. Tourtellotte ME, Jacobs RE. PHYSIOLOGICAL AND SEROLOGIC COMPARISONS OF PPLO FROM VARIOUS SOURCES. Ann N Y Acad Sci. 1960; doi:10.1111/j.1749-6632.1960.tb42718.x

27. Hahn RG, Kenny GE. Differences in arginine requirement for growth among arginine utilizing Mycoplasma species. J Bacteriol. 1974;

28. Levina G, Barkhatova O, Rakovaskaya I. Stress causes formation of morphologically new type of Mycoplasma hominis. FEBS OPEN Bio. p. 144.

29. Maniloff J, Nealson K, Psenner R, Maria L, Robert F. Nannobacteria?: Size Limits and Evidence. Science (80-). 1997;

30. Olavi Kajander E, Ciftçioglu N, Seegmiller JE. Nanobacteria: An alternative mechanism for pathogenic intra-and extracellular calcification and stone formation. Med Sci. 1998; doi:10.1073/pnas.95.14.8274

31. Keren I, Kaldalu N, Spoering A, Wang Y, Lewis K. Persister cells and tolerance to antimicrobials. FEMS Microbiol Lett. 2004; doi:10.1016/S0378-1097(03)00856-5

32. Kussell E, Leibler S. Phenotypic diversity, population growth, and information in fluctuating environments. Science. 2005; doi:10.1126/science.1114383

33. Lewis K. Persister Cells. Annu Rev Microbiol. 2010; doi:10.1146/annurev.micro.112408.134306

34. Proctor RA, Kriegeskorte A, Kahl BC, Becker K, LÃ¶ffler B, Peters G. Staphylococcus aureus Small Colony Variants (SCVs): a road map for the metabolic pathways involved in persistent infections. Front Cell Infect Microbiol. 2014; doi:10.3389/fcimb.2014.00099

35. Edwards AM. Phenotype switching is a natural consequence of Staphylococcus aureus replication. J Bacteriol. 2012; doi:10.1128/JB.00948-12

36. Kriegeskorte A, Lorè NI, Bragonzi A, Riva C, Kelkenberg M, Becker K, et al. Thymidine-dependent Staphylococcus aureus small-colony variants are induced by trimethoprim-sulfamethoxazole (SXT) and have increased fitness during SXT challenge. Antimicrob Agents Chemother. 2015; doi:10.1128/AAC.00742-15

37. Painter KL, Strange E, Parkhill J, Bamford KB, Armstrong-James D, Edwards AM. Staphylococcus aureus adapts to oxidative stress by producing H2O2-resistant small-colony variants via the SOS response. Infect Immun. 2015; doi:10.1128/IAI.03016-14

38. Proctor RA, Van Langevelde P, Kristjansson M, Maslow JN, Arbeit RD. Persistent and relapsing infections associated with small-colony variants of staphylococcus aureus. Clin Infect Dis. 1995; doi:10.1093/clinids/20.1.95

39. Chernov VM, Chernova OA, Mouzykantov AA, Ponomareva AA, Trushin M V., Gorshkov O V., et al. Phytopathogenicity of avian mycoplasma Mycoplasma gallisepticum S6: Morphologic and ultracytostructural changes in plants infected with the vegetative forms and the viable but nonculturable forms of the bacterium. Microbiol Res. 2010; doi:10.1016/j.micres.2009.07.001

40. Medvedeva ES, Davydova MN, Mouzykantov AA, Baranova NB, Grigoreva TY, Siniagina MN, et al. Genomic and proteomic profiles of Acholeplasma laidlawii strains differing in sensitivity to ciprofloxacin. Dokl Biochem Biophys. 2016; doi:10.1134/S1607672916010075

41. Lysnyansky I, Ayling RD. Mycoplasma bovis: Mechanisms of resistance and trends in antimicrobial susceptibility. Frontiers in Microbiology. 2016. doi:10.3389/fmicb.2016.00595

42. Ito S, Shimada Y, Yamaguchi Y, Yasuda M, Yokoi S, Ito SI, et al. Selection of Mycoplasma genitalium strains harbouring macrolide resistance-associated 23S rRNA mutations by treatment with a single 1 g dose of azithromycin. Sex Transm Infect. 2011; doi:10.1136/sextrans-2011-050035

43. Dégrange S, Renaudin H, Charron A, Pereyre S, Bébéar C, Bébéar CM. Reduced susceptibility to tetracyclines is associated in vitro with the presence of 16S rRNA mutations in Mycoplasma hominis and Mycoplasma pneumoniae. Journal of Antimicrobial Chemotherapy. 2008. doi:10.1093/jac/dkn118

44. Amram E, Mikula I, Schnee C, Ayling RD, Nicholas RAJ, Rosales RS, et al. 16S rRNA gene mutations associated with decreased susceptibility to tetracycline in Mycoplasma bovis. Antimicrob Agents Chemother. 2015; doi:10.1128/AAC.03876-14

45. Balwit JM, Langevelde P Van, Vann JM, Proctorn RA. Gentamicin-resistant menadione and hemin auxotrophic staphylococcus aureus persist within cultured endothelial cells. J Infect Dis. 1994; doi:10.1093/infdis/170.4.1033

